# Screening and structural characterization of a nanobody targeting the thermostable green fluorescent protein

**DOI:** 10.64898/2025.12.23.696179

**Authors:** Yaning Li, Weijie Gu, Aimin Ni, Dianfan Li, Tingting Li

## Abstract

Nanobody–fluorescent protein pairs are powerful tools in imaging, protein purification, and structural biology studies. While thermostable GFP (TGP) offers improved characteristics over conventional GFP, nanobodies that specifically recognize TGP remain relatively limited. Here, we report the screening and identification of a synthetic nanobody, Sb32, that binds TGP with nanomolar affinity using the ribosome technique. The crystal structure of the Sb32–TGP complex, solved at 1.82 Å resolution, revealed an unusual binding mode in which Complementarity Determining Region 2 (CDR2) provides the major contribution, rather than the typically dominant CDR3. Moreover, the interface is stabilized by an extensive hydration network, which compensates for relatively few direct contacts and may explain the fast association and dissociation kinetics observed for this complex. These findings expand the repertoire of TGP-specific nanobodies and highlight an alternative strategy by which nanobodies achieve high affinity. Sb32 provides a new reagent for TGP-based applications, with potential utility in membrane protein purification and structural studies.

## Introduction

Llamas and sharks produce heavy chain–only antibodies that lack the light chains and the constant regions present in conventional IgG antibodies. The <u>v</u>ariable heavy domain of <u>h</u>eavy chain (VHH), commonly known as a nanobody due to its small size (∼14 kDa), has emerged as a powerful tool for both basic and biomedical applications. Caplacizumab, the first Food and Drug Administration (FDA)-approved nanobody drug used to treat adult patients with acquired thrombotic thrombocytopenic purpura ^1^, marked a milestone in this field and has spurred further basic and clinical research into their development and application in cancer, ^2–8^ autoimmune immunotherapy, ^9–11^ and infectious diseases ^12^.

The many advantages of nanobodies stem from their single-domain nature and small size. At the organism or tissue level, their compactness allows high permeability, rapid diffusion, and fast clearance. This makes nanobodies attractive for in vivo imaging applications ^13^, such as radiolabeled positron emission tomography (PET) in oncology ^14^. At the molecular level, their small size enables recognition of epitopes that are otherwise inaccessible to conventional IgG antibodies, establishing them as promising neutralizing agents against bacterial ^15^, fungal ^16^, parasitic, and viral ^15,17–21^ pathogens.

The single-domain architecture of nanobodies also contributes to exceptional stability, withstanding heat and chemical stresses such as reducing agents. This property facilitates long-term storage, transport, production ^22^, and intracellular applications, where the reducing environment often disrupts disulfide bonds of conventional antibodies. Such cellular stability has enabled the development of innovative tools, exemplified by the recently described ALFA Nb-guided endogenous labeling (ANGEL) strategy ^23^, where fluorescently labeled nanobodies are engineered to selectively report successful genome edits and enable dynamic protein studies in living cells. Further, the remarkable stability of nanobodies not only supports their systemic administration but also makes inhaled delivery feasible as nanobodies can remain effective after nebulization ^24^. This route of administration is especially attractive for combating respiratory viral infections, as it allows direct targeting of the respiratory tract while bypassing the need for intravenous infusion or hospital-based care. Such an approach could significantly reduce the burden on healthcare facilities, particularly during widespread and prolonged outbreaks such as SARS-CoV-2, when medical resources are strained and timely access to treatment is critical.

Nanobodies have also transformed structural biology, particularly in the membrane protein field. Tightly bound nanobodies can serve as crystallization chaperones, addressing the challenge that membrane proteins often lack suitable surfaces for crystal packing ^25–28^. In cryo-electron microscopy (cryo-EM), nanobodies can act as fiducial markers to improve particle alignment ^29^. Their modularity and ease of engineering have further expanded their utility, giving rise to variants such as macrobodies (nanobody fused rigidly to maltose-binding protein, MBP) ^30^, megabodies (chimeras of nanobodies with larger scaffolds) ^31,32^, NabFabs (anti-nanobody Fabs) ^33^, Legobodies (MBP variants noncovalently linked to NabFabs) ^34–36^, and covalently linked homo- or hetero Di-Gembodies ^37^. These engineered constructs increase the apparent size of small proteins, helping push the boundaries of current molecular weight limits accessible for the cryo-EM method. Furthermore, unlike IgG/Fab fragments, which often contain crevices suited for linear epitopes, nanobodies preferentially recognize three-dimensional conformational epitopes as such crevices are difficult to form within their single-domain structures. This specificity can be exploited to stabilize dynamic protein conformations that are otherwise difficult to capture ^38^. Prominent examples include nanobodies that lock G protein–coupled receptors (GPCRs) in active/inactive states for signaling studies ^38–41^, and trap transporters ^42,43^ in specific snapshots for structure determination.

The thermostable green fluorescent protein (TGP) is a heat-resistant GFP originated from coral ^44^. As a bright fusion partner, TGP facilitates convenient monitoring of membrane protein expression, solubilization, and purification ^45–48^, similar to conventional GFP. Compared with the jellyfish GFP, TGP offers several advantages as a fusion partner, including higher expression yields of target protein and more accurate reporting of membrane protein stability ^49,50^ in fluorescence-detection size exclusion chromatography (FSEC) assays ^51^. However, unlike the jellyfish GFP, nanobodies against TGP remain scarce ^49,52^, limiting their broader use as affinity reagents or as structural biology tools. For example, a recent study demonstrated the use of a nanobody–GFP complex to construct a ‘Di-Gembody’, in which the anti-GFP nanobody is covalently linked to another nanobody targeting a membrane protein of interest ^37^. By increasing the apparent size of the target protein by ∼40 kDa, this approach facilitates cryo-EM analysis of small membrane proteins that are otherwise refractory for successful image alignments. Being more stable than the conventional GFP, TGP may provide a better alternative in such applications. However, the DiGembody strategy often requires trial-and-error tests because the nanobody–fluorescence protein complex may sterically clash with the target, depending on the binding mode ^37^. Therefore, having multiple nanobody–fluorescence protein options is desirable for screening in such applications.

In this study, we report the screening and structural characterization of a high-affinity synthetic nanobody (sybody) against TGP. The crystal structure solved at 1.82 Å resolution reveals the molecular details of sybody–TGP interactions and provides insight into the binding kinetics. This newly characterized sybody expands the toolkit for membrane protein purification and structural studies.

## Results

### Screening of nanobody against TGP from a synthetic library

To identify nanobodies specific for TGP, we performed ribosome display using a synthetic concave library (see Methods) ^53^. Briefly, a high-quality synthetic mRNA library was displayed on ribosomes in a test tube. mRNA encoding potential TGP binders was captured using beads coated with biotinylated TGP and reverse-transcribed into cDNA. The number of cDNA copies was semi-quantified by quantitative polymerase chain reaction (qPCR) using a standard curve from calibration experiments with known amounts of plasmid DNA. The copy number was determined to be 2.8 × 10^8^ (**Fig. 1a**), a diversity that was within the range of electroporation transformation efficiency. The cDNA was then subcloned into a phage display library for further panning. During the panning process, phage particles were first bound with beads immobilized with biotinylated TGP, washed, before being eluted by tryptic digestion. The copy number of phage particles were also semi-quantified by qPCR. The same process was repeated using a maltose binding protein (MBP) as a control to assess the enrichment of TGP-specific binders.

**Figure 1.**
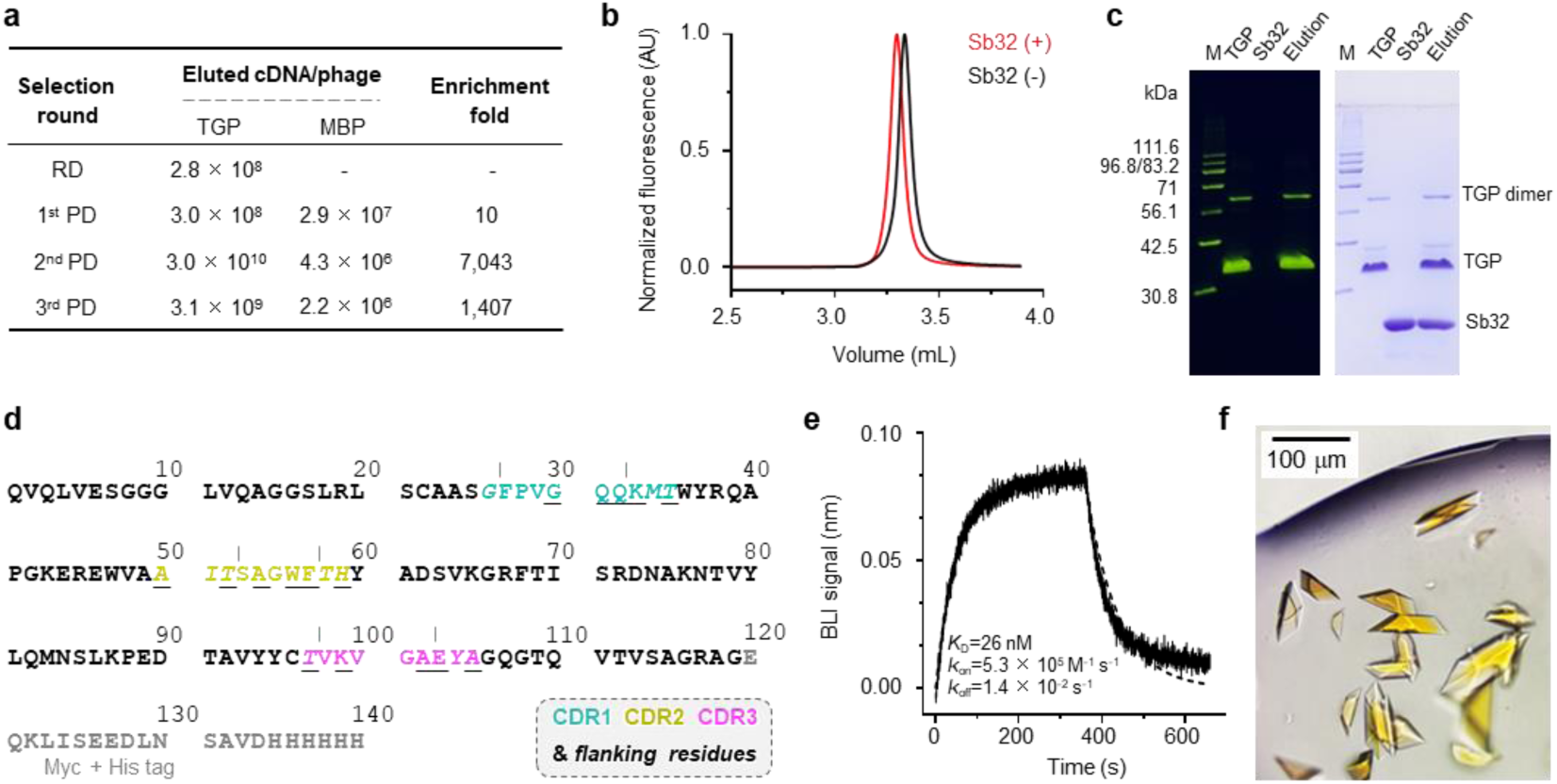
Screening and biophysical characterization of the Sb32-TGP complex. **a** Quality assessment of the nanobody selection steps. **b** Fluorescence-based size exclusion chromatography of TGP in the absence (black) or presence (red) of Sb32. **c** SDS-PAGE of TGP, Sb32, or the co-eluted peak fraction (Elution). The gel was visualized by in-gel fluorescence (left) or Coomassie blue staining (right). The apparent molecular weights of home-made fluorescence markers (M) were indicated on the left. **d** Sequence of Sb32. Amino acids in the CDRs (marked by vertical lines) and the CDR-flanking residues (italicized), are color-coded as indicated in the dashed box. Randomized residues are underlined. Tag residues are shown in grey. **e** Kinetic profile of Sb32 binding to TGP. A sensor coated with streptavidin was saturated with 2 μg mL^-1^ of biotinylated TGP. The sensor was then soaked in 10 nM of Sb32 before back-soaked with sybody-free buffer at indicated time points. Data (solid line) were fitted with the built-in software (dotted line) from which the kinetic parameters were calculated. **f** Crystals from crystallization trials of the Sb32-TGP complex. The precipitant solution contained 0.2 M ammonium sulfate, 20% (w/v) polyethylene glycol 8,000, and 0.1 M Na-MES pH 6.5. Crystals were grown in a sitting drop plate in an incubator at 20 °C.

TGP-specific particles were enriched by 10 folds after the first phage display cycle, and the enrichment fold increased drastically to over 7,000 fold after the second round of phage display (**Fig. 1a**). To screen high-affinity binders, a more stringent panning condition was employed for the third phage-display cycle (see Methods). The resulting library was subcloned into an expression vector and the periplasmic extract from clones in a 96-well plate was used to screen binders using fluorescence-detection size-exclusion chromatography (FSEC). We chose a positive clone named Sb32 for downstream characterization.

Sb32 caused a leftward shift of V_e_ of TGP on FSEC (**Fig. 1b**), indicating formation of a complex. Preparative gel filtration and SDS-PAGE analysis (**Fig. 1c**) confirmed the co-elution of Sb32 and TGP. The deduced amino acid sequence of Sb32 (**Fig. 1d**) was different than the two previously characterized sybodies, Sb44 ^49^ and Sb92 ^52^. Biolayer interferometry (BLI) analysis showed a *k*_on_ of 5.3×10^5^ M^-1^ s^-1^ and a *k*_off_ of 1.4×10^-2^ s^-1^ (**Fig. 1e**). The resulting equilibrium dissociation constant (*K*_D_) was 26 nM, indicating the formation of a relatively tight complex.

### Crystal structure determination of the Sb32-TGP complex

To gain structural insight into Sb32–TGP interactions, the complex was purified and crystallized. Yellow crystals formed readily in precipitant solutions containing ammonium sulfate and high–molecular weight polyethylene glycol (**Fig. 1f**). Crystals typically appeared within one day and grew to a final size of ∼100 μm × 30 μm × 10 μm within ten days at room temperature (20 °C).

The crystals diffracted to 1.82 Å resolution at a synchrotron light source. Data processing revealed that the crystals belonged to the *C121* space group with unit cell dimensions a = 162.47, b = 89.92, c = 111.24 Å, and β = 112.12°. Matthews coefficient analysis suggested either three complexes per asymmetric unit (59.2% solvent content) or four complexes (45.6% solvent content). Molecular replacement using individual TGP and sybody models identified three copies of the Sb32–TGP complex in the asymmetric unit. Structural models were built in Coot and refined with Phenix, yielding *R*_work_ and *R*_free_ values of 15.67% and 17.28%, respectively. Despite the absence of non-crystallographic symmetry (NCS) restraints during refinement, the Cα root-mean-square deviation (RMSD) among NCS-related molecules was only 0.12 Å. Therefore, a single representative pair-Chain A (TGP) and Chain B (Sb32) - was selected for subsequent structural descriptions.

### Overall structure of the complex

Sb32 binds TGP at the side of the β-barrel, burying 529.7 Å^2^ of surface area (**Fig. 2a**). Consistent with the notion that nanobodies generally recognize three-dimensional epitopes, Sb32 engages with TGP via residues distributed across strands β1, β5, β6, and the flanking regions of β4 and β9 (**Fig. 2b**). All three complementarity-determining regions (CDRs) contribute to TGP binding. However, unlike most naturally occurring nanobodies, where CDR3 provides the major contribution, Sb32 interacts with TGP primarily through CDR2. This binding mode reflects the design of the Concave library ^53^: CDR3 was intentionally kept very short, with only five residues, and only two of these residues, facing away from the nanobody core, were randomized (**Fig. 1d**). As described in the original paper ^53^, this design, together with additional residues forming a stable hydrophobic core, maintains a concave shape of the antigen-recognition surface. To compensate for the limited diversity of CDR3, several residues in CDR1, CDR2, and their flanking regions were randomized (**Fig. 1d**), effectively shifting the paratope toward CDR2 and explaining why CDR2 makes the predominant contribution to TGP binding.

**Figure 2.**
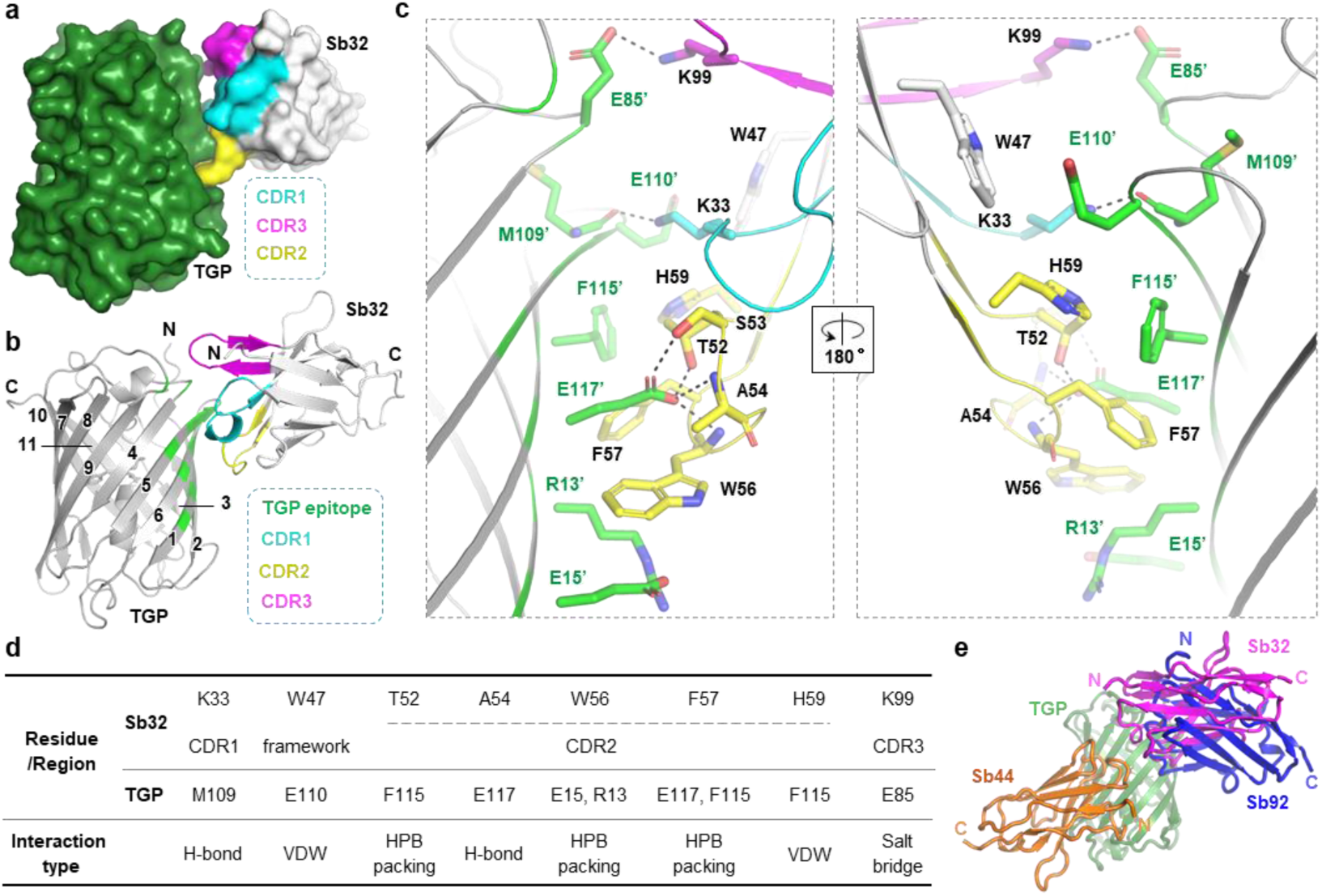
Structure of the Sb32-TGP complex. **a** Surface representation of Sb32 (light grey) bound with TGP (dark green). CDR1, CDR2, and CDR3 and their flanking residues are colored cyan, yellow, and magenta, respectively. **b** Cartoon representation of Sb32 (light grey) bound with TGP (dark green). CDR1, CDR2, and CDR3 and their flanking residues are colored cyan, yellow, and magenta, respectively. TGP epitope residues are colored green. TGP β-strands are marked with numbers. The N- and C-termini of Sb32 and TGP are labeled with N/C (**ii**). **c** The Sb32-TGP interaction network. TGP residues (green) and Sb32 residues (colored coded by CDRs as indicated in panels **a/b**) at the interaction interface are shown as sticks. Mainchain atoms are omitted for stick representation unless they are involved in direct interactions. Framework residues are colored grey. Hydrogen bonds and salt bridges with distances within 3.4 Å are shown as black dashed lines. **d** Summary of Sb32-TGP interaction. H-bond, hydrogen bonding; HPB, hydrophobic; VDW, Van der Waals. **e** Comparison of the binding mode between Sb32 (this work) and two previously reported TGP-targeting sybodies, Sb44 (PDB ID 6LZ2)^49^ and Sb92 (PDB ID 7CZ0)^52^. Structure models of Sb44 (orange) and Sb92 (blue) were superposed onto the Sb32-TGP complex with TGP (green) as the reference.

### Molecular basis for Sb32-TGP binding

Shape complementarity is a key determinant in protein-protein interactions ^54^. In the Sb32–TGP complex, this complementarity is exemplified by the insertion of the bulky Trp56 of CDR2 into a surface dent formed by the aliphatic portions of three charged TGP residues: Glu117’, Glu15’, and Arg13’ (a prime denotes TGP residues hereafter) (**Fig. 2a**, **2c**, **2d**). Glu117’ plays a central role in the interaction, as Sb32 forms four of its five total hydrogen bonds with this residue in a tetradentate arrangement, by interacting with the side-chain hydroxyls of Thr52 and Ser53, and the backbone nitrogen atoms of Ala54 and Trp56 (**Fig. 2c, 2d**). At the ‘base’ of the dent, an intramolecular salt bridge between Glu15’ and Arg13’ further stabilizes the interaction network.

Interactions near CDR2 are reinforced by hydrophobic packing involving Sb32 Phe57 and TGP Phe115’, as well as van der Waals contacts between His59 of Sb32 from Sb32 and Phe115’/Glu110’ from TGP. In CDR1, Lys33 forms a hydrogen bond with the backbone of TGP Met109’. In CDR3, Lys99 establishes the sole intermolecular salt bridge with TGP Glu85’. Beyond the CDRs, a framework residue, Trp47, packs closely against Glu110’, contributing to additional complex stability (**Fig. 2c, 2d**).

Superimposing the structures of two previously known TGP-targeting sybodies, Sb44 and Sb92, onto the Sb32-TGP complex (**Fig. 2e**) reveals that the Sb32 and Sb92 epitopes partially overlap. Both adopt a head-to-head configuration with Sb44. Notably, the C-terminus of Sb32 points in a different direction from that of Sb92, making Sb32 an alternative partner for Di-Gembody assembly^37^ should Sb92 or Sb44 introduce steric hindrance in such applications.

### A hydration interaction network

We observed a notable gap between CDR1/CDR3 and TGP (**Fig. 2a**), which is unusual given the nanomolar affinity of the Sb32–TGP complex. A closer inspection revealed an extensive hydration network at the interface (**Fig. 3a, 3b**). Consistent with the observation that CDR2 forms the primary direct contacts with TGP, this region is only modestly solvated. CDR3 is located at the more peripheral space of the complex that is more open to the bulk solvent; as expected, this region also contained fewer ordered water molecules (**Fig. 3b, 3c**). By contrast, several highly coordinated water molecules were observed around CDR2 (**Fig. 3b, 3c**).

**Figure 3.**
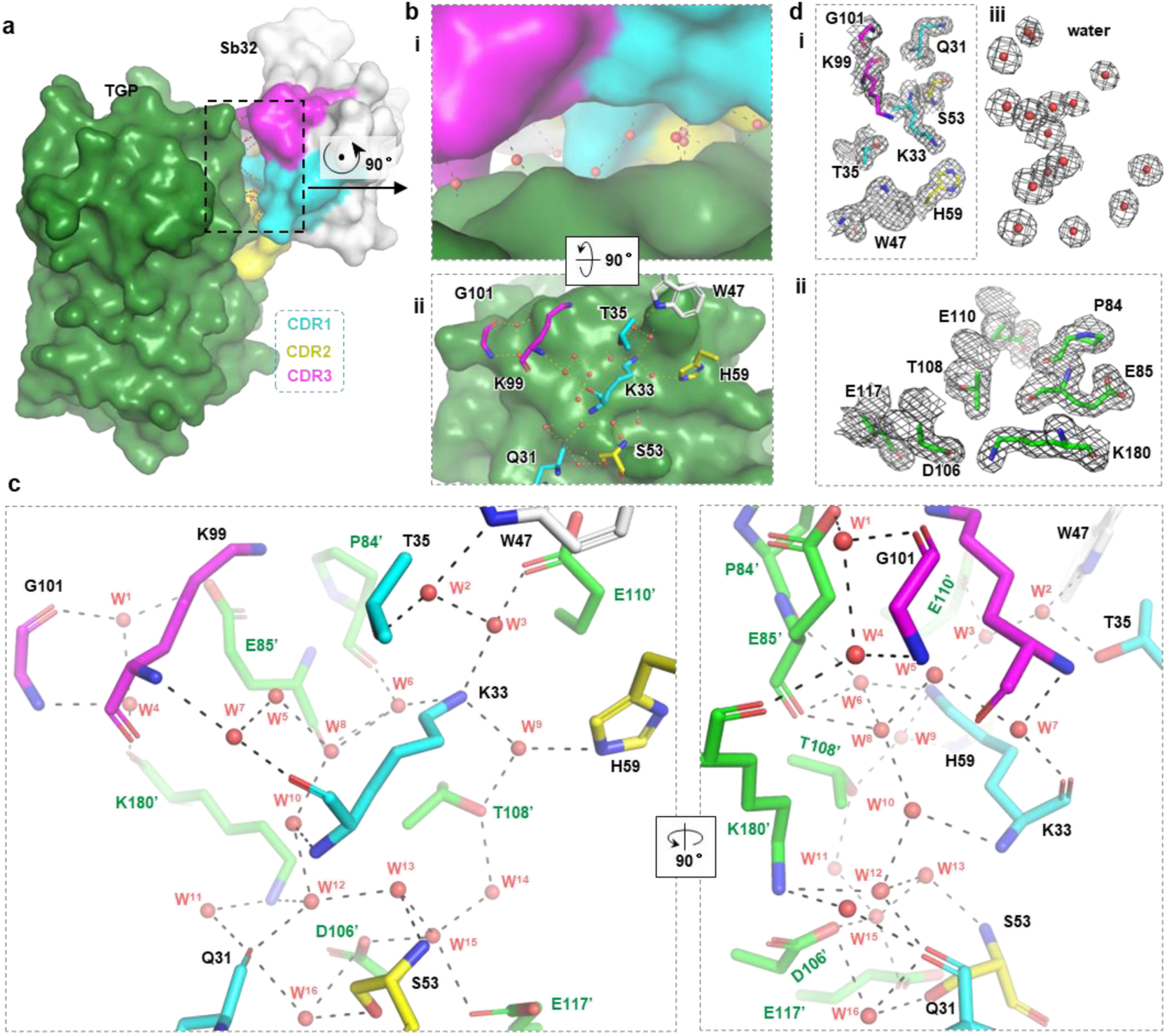
The hydration interaction network that stabilizes the Sb32-TGP complex. **a** Surface representation of Sb32 (light grey) bound with TGP (dark green). CDR and their flanking residues are color-coded as indicated in the dashed box at the right bottom. The black dashed box indicates the gap between Sb32 and TGP where the lack of direct contacts was observed. **b** An expanded view of the dashed box in **a**. Illustration was made to contain surface representations for proteins (**i**) or a mix of surface representation (TGP) and stick representation (Sb32) (**ii**). Water molecules are shown as red spheres. Only the Sb32 residues are labeled. **c** Detailed interaction network at the hydrated interface. Black, green, and red letters mark protein residues from Sb32 (cyan for CDR1, yellow for CDR2, and magenta for CDR3), TGP (green sticks), and water molecules (red sphere) respectively. **d** Electron density map (2*F*_o_-*F*_c_) (grey mesh) of key residues from Sb32 (i) and TGP (ii), as well as water molecules discussed in the text contoured at a σ level of 1.0.

Overall, 16 ordered water molecules were identified at the Sb32–TGP interface, mediating interactions without direct atomic contact. Specifically, Water 1 and 4 bridge the backbone of Gly101 from Sb32 with two charged TGP residues, Glu85’ and Lys180’. Water 2 and 3 connect the negatively charged Glu110’ of TGP with Lys33 of Sb32, and also interact with the hydroxyl of Thr35 and the indole nitrogen of Trp47. Waters 5, 7, and 8 form a network linking the main chains of Lys33 and Lys99 from Sb32 with Glu85’ from TGP, while Water 6 mediates interactions between the carbonyls of Pro84’/Glu85’ from TGP and Lys33 from Sb32. Water 9 is coordinated by the side chains of His59 and Lys33 from Sb32 and Thr108’ from TGP. Waters 8, 10, and 12 bridge backbone atoms of Sb32 Lys33 with Glu85’ and Lys180’ from TGP. Waters 11 and 12 connect the backbone of Sb32 Gln31 with the amine group of Lys180’ in TGP. Waters 13, 14, and 15 form a hydrogen-bond relay linking the side-chain oxygen atoms of TGP Thr108’ and Asp106’ to the backbone nitrogen of Sb32 Ser53. Finally, Water 16 coordinates with TGP Asp106’ and Sb32 residues Gln31 and Ser53 (**Fig. 3c, 3d**).

## Discussion

Nanobodies are increasingly recognized as versatile tools in basic research. In particular, nanobody–fluorescent protein pairs, such as those targeting GFP or mCherry, are used to monitor pathogen–host interactions ^55^, track cellular events ^56^, facilitate affinity purification of fusion membrane proteins ^57^, and aid cryo-EM structural determination ^37^. As a thermostable fluorescent protein, TGP provides an improved alternative for biological studies, yet structurally characterized nanobodies against TGP have been lacking. In this study, we filled this gap by generating a high-affinity sybody against TGP and revealing high-resolution details of the antigen–antibody interactions. These structural insights provide a framework for rational engineering of the complex, both components of which are amenable to genetic and chemical modifications.

The binding kinetics of Sb32–TGP are noteworthy. Despite a fast dissociation rate, the overall affinity is reported in the nanomolar range, reflecting a rapid association rate. In applications where higher apparent affinity or slower dissociation is desired, divalent forms of Sb32 could be employed to leverage avidity effects. Based on our previous experience, direct divalent fusions or Fc-mediated dimerization can increase apparent affinity by up to three orders of magnitude.

Interestingly, despite its nanomolar affinity, the footprint of Sb32 on TGP is relatively small, with a buried surface area of only 519 Å^2^, which is substantially lower than those observed from our previous nanomolar-affinity sybodies ^18–20,58^. Specifically, Sb44 and Sb92 engage with TGP with a buried surface area of 836 Å^2^ and 619 Å^2^, respectively^49,52^. In line with this, Sb44 (*K*_D_ = 4.4 nM)^49^ and Sb92 (*K*_D_ = 3.3 nM)^49^ bind TGP with *K*_D_ values approximately one order of magnitude lower than Sb32 (*K*_D_ = 26 nM). The limited direct contact in Sb32-TGP interaction appears to be compensated by an extensive network of ordered water molecules at the interface. Such hydration-mediated interactions may also explain the fast-on/fast-off kinetic behavior of the Sb32–TGP complex: water-mediated interactions are more dynamic than direct contacts, as interface water molecules, unlike protein residues, would readily exchange with the bulk solvent. This property suggests that the affinity of this type of complex may be more sensitive to environmental factors, which in cellular contexts could provide new opportunities for modulating protein–protein interactions. Further structural characterization of similar hydration-mediated interfaces could inform de novo antibody design and engineering algorithms, advancing our understanding of protein interactions and expanding the toolkit for molecular biology research.

## Methods

### Purification of TGP

TGP was expressed in *Escherichia coli* as a C-terminally hexa-histidine-tagged protein. BL21 (DE3) cells carrying the plasmid pETSG^59,60^ was induced at an optical density at 600 nm (OD_600_) of 0.6–0.8 for 20 h at 20 °C using 1 mM isopropyl-β-_D_-thiogalactopyranoside (IPTG). Cells from 100 mL culture were suspended in Buffer A (150 mM NaCl, 50 mM HEPES pH 7.5) supplemented with 1 mM PMSF. Cells were disrupted with an ultrasonic cell disruptor (Cat. JY92-IIN, SCIENTZ, CN) for 10 min in an ice-water bath. The cell lysate was clarified by centrifugation at 20,000 × g for 15 min at 4 °C. The supernatant was heated at 65 °C for 15 min to denature non-target proteins, cooled in an ice-water bath, followed by centrifugation at 20,000 × g for 15 min at 4 °C to remove precipitates. The supernatant containing heat-resistant TGP was incubated with 0.6 mL of Ni-NTA slurry for 1 h with gentle agitation at 4 °C. The beads were loaded into a gravity column and washed with 20 column volume (CV) of Buffer A supplemented with 30 mM imidazole. TGP was then eluted using 400 mM imidazole in Buffer A. TGP was quantified using absorbance at 280 nm measured on a Nanodrop machine with the theoretical molar extinction coefficient of 31,985 M^-1^ cm^-1^.

### Purification of sybody

Sybody was expressed in *E. coli* as a C-terminally His-tagged proteins. MC1061 cells carrying the plasmid pSB_init_Sb32 was induced at OD_600_ of 0.6–0.8 for 20 h at 22 °C. Cells were lysed by osmotic burst as follows. Cells from 1 L of culture was resuspended in 20 mL high-osmolarity TES buffer (0.5 M sucrose, 0.5 mM EDTA, and 0.2 M Tris-HCl pH 8.0) for dehydration at 4 °C for 0.5 h. Dehydrated cells were rehydrated abruptly by diluting the cells with 40 mL of ice-cold MilliQ H_2_O at 4 °C for 1 h. Periplasmic extracts were collected as the supernatant fraction after centrifugation at 20,000 × g at 4 °C for 30 min. After being adjusted to contain 150 mM of NaCl, 2 mM of MgCl_2_, and 20 mM of imidazole, the supernatant was added with two milliliters of Ni-NTA resin pre-equilibrated with 20 mM imidazole in Buffer B (150 mM NaCl and 20 mM HEPES pH 7.5). Batch binding was performed by gentle mixing the two at 4 °C for 1.5 h. The resin was then packed into a gravity column and washed with 20 CV of 30 mM imidazole in Buffer B. The sybody was eluted using 250 mM imidazole in Buffer B. Sybody proteins were quantified using absorbance at 280 nm measured on a Nanodrop machine with a molar extinction coefficient of 25,565 M^-1^ cm^-1^.

### Sybody selection—ribosome display and phage display

TGP containing a C-terminal Avi-tag (GLNDIFEAQKIEWHE) (TGP_Avi_) and a 3C-cleavable His-tag ^60^ was purified the same way as described above. For enzymatic biotinylation, the protein at 4.5 mg mL^−1^ was incubated with 0.13 mg mL^−1^ of His-tagged BirA, 0.72 mg mL^−1^ His-tagged 3C protease, 5 mM ATP, 10 mM magnesium acetate, 0.23 mM biotin in 500 μL volume and incubated at 4°C for 11 h. The reaction mixture was mixed with 0.5 mL pre-equilibrated Ni-NTA resin for 1.5 h to remove His-tagged 3C protease and BirA. The flowthrough fraction containing biotinylated TGP_Avi_ was further purified by gel filtration on a Superdex 200 increase 10/300 GL column pre-equilibrated with buffer containing 150 mM NaCl, 20 mM Tris-HCl pH 8.0. Fractions with absorbance at 493 nm were pooled, adjusted to 0.6 mg mL^−1^, aliquoted, flash-frozen with liquid nitrogen, and stored at -80°C before use.

In vitro translation was performed using the PUREfrex 2.1 kit (Cat. PF213-0.25-EX, Genefrontier, Chiba, Japan) supplemented with the disulfide bond isomerase DsbC (DS supplement, Cat. PF005-0.5-EX, Genefrontier) per the manufacturer’s instructions and as described previously ^60^. The in vitro translation products were diluted with 100 μL ice-cold buffer containing 150 mM NaCl, 50 mM magnesium acetate, 0.5%(w/v) BSA, 0.1%(w/v) Tween 20, 0.5%(w/v) heparin, 1 μL RNaseIn (RNase inhibitor), and 50 mM Tris acetate pH 7.4. After centrifugation at 20,000 × g for 5 min at 4 °C, biotinylated TGP_Avi_ was incubated with the supernatant for 20 min.

Pull-down was then performed by adding streptavidin (Dynabeads Myone Streptavidin T1) beads into the mixture to recover mRNA encoding TGP binders. After reverse transcription with the primer 5′-CTTCAGTTGCCGCTTTCTTTCTTG-3′, the cDNA library was amplified using primer pairs 5′-ATATGCTCTTCTAGTCAGGTTCAGCTGGTTGAGAGCG-3′ and 5′-TATAGCTCTTCATGCGCTCACAGTCACTTGGGTACC-3′. The DNA products were gel-purified, digested with BspQI, and ligated into the vector pDX_init digested with the same enzyme. Ligation products were transformed into 0.35 mL of *E.coli* SS320 competent cells by electroporation using a Bio-Rad MicroPulser Electroporator with a voltage of 2,400 V. Phage display was performed in a 96-well plate coated with 60 nM neutravidin (Cat. 31000, Thermo Fisher Scientific, Waltham, MA, USA). To recover phage, 40 mL of cell culture were mixed with 10 mL ice-cold solution containing 20 % PEG 6000 and 2.5 M NaCl to precipitate the phage particles. After a 30-min incubation, the mixture was centrifuged at 3,220 × g for 30 min at 4 °C. Phage particles in the pellet fraction were resuspended in 1 mL of PBS buffer. Phage particles were clarified by another centrifugation step at 20,000 × g for 30 min at 4 °C. The supernatant was diluted into 5.0 mL of panning solution (0.5%(w/v) BSA, 0.1%(w/v) Tween 20, 150 mM NaCl, 50 mM Tris-HCl pH 7.4) and then incubated with 50 nM biotinylated TGP_Avi_ for 10 min. The mixture (4.7 mL) was then added into 47 wells of an aforementioned neutravidin-coated 96-well plate for incubation. After 10 min, the plate was washed, and phage particles were proteolytically eluted from the beads by adding trypsin to a final concentration of 0.25 mg/mL. The enriched phage particles (4.7 mL) were used to infect 45 mL of *E.coli* SS320 cells at an OD_600_ of 0.6. After culturing at 37 °C overnight, 1 mL of culture was seeded into 50 mL of fresh medium and the cells were allowed to grow to OD_600_ of 0.6. Cells (10 mL) were then infected with M13KO7 helper phage to assemble the whole phage particles. Phage particles were harvested as described above and resuspended in 1 mL of PBS. A hundred microliter of panning solution containing 5 × 10^11^ phage particles were used to perform the second and third round phage display using 12 μL MyOne Streptavidin C1 Dynabeads as the immobilizing matrix. TGP_Avi_ was included at 50 nM and 5 nM for the second and third phage display, respectively. Phagemid from the 3^rd^ phage display were extracted and subcloned into pSb_init vector using fragment-exchange (FX) cloning and transformed into *E. coli* MC1061 for expression and purification.

MC1061 colonies were cultured in a 96-well plate. After overnight induction, cells were spun down and treated by osmotic shock as mentioned above. For FSEC screening, 50 μL of periplasmic extraction were incubated with 1 μL of TGP to have a final TGP concentration of 3-5 μg mL^-1^. One microliter of the mixture was applied onto an analytic gel filtration column connected in a Shimadzu HPLC machine equipped with a fluorescence detector (RF-20A, Shimadzu). The elution profile was monitored by fluorescence with an excitation wavelength of 482 nm and an emission wavelength of 508 nm.

### Biolayer interferometry assay

Biolayer interferometry assay was performed using an Octet system. Biotinylated TGP_Avi_ was immobilized onto a streptavidin SA sensor (ForteBio, Cat 18-5019) by incubating the sensor in TGP_Avi_ (2 μg mL^−1^) solution in a buffer containing 0.005%(v/v) Tween20, 150 mM NaCl, and 20 mM HEPES pH 7.5. The sensor was then placed in a sybody-free buffer until the signal became flat. The SA sensor was incubated with the same buffer supplemented with 10 nM sybody (Association phase). The data was fitted using a 1:1 stoichiometry with the built-in software Data Analysis 10.0.

### Crystallization

The TGP protein used for crystallization contained residues 1-218 of TGP with a SGGGSGGG linker between the C-terminal octa-histidine tag ^60^. TGP and Sb32 were mixed at a molar ratio of 1:1.2, and the complex was purified on a gel filtration column in a running buffer containing 150 mM NaCl, 20 mM HEPES pH 7.5. Pooled fractions with absorbance at 493 nm were concentrated to 20 mg mL^-1^ using a 10-kDa cut-off filtration membrane (Cat. UFC501096, Merck Millipore, Burlington, MA, USA). Sitting drop crystallization was performed using a Gryphon robot by depositing 0.15 μL of the precipitant solution on top of equal volume of protein with 70 μL of solutions in the reservoir. Crystallization plates were incubated at 20 °C in an incubator. Crystals grew in a precipitant solution containing 0.2 M ammonium sulfate; 20% (w/v) polyethylene glycol 8,000, and 0.1 M Na-MES pH 6.5.

### **X-** ray Diffraction data collection

Crystals were harvested on day 21 using a MiTeGen loop (Cat. M5-L18SP series, Ithaca, NY, USA) under an Olympus light microscope (Model BX43F, Olympus, Tokyo, Japan). Sigle crystals harvested on loops were step-wise soaked in the precipitant solution supplemented with 15 %(v/v) glycerol before being plunged into liquid nitrogen for cryo-cooling. X-ray diffraction data were collected at beamlines 18U1 at the National Facility for Protein Science in Shanghai (NFPS) at Shanghai Synchrotron Radiation Facility with a 50 × 50 μm beam on a Pilatus 6M detector with oscillation of 0.5° and a wavelength of 0.97930 Å. Diffraction spots were integrated using XDS ^61^, scaled and merged using Aimless ^62^. The TGP-Sb32 structure was solved by molecular replacement using Phaser ^63^ with three TGP monomer and three sybody molecules taken from the Sb44-TGP structure (PDB ID 6LZ2) ^60^ as the searching models. The model was built with 2*F_o_*-*F_c_* maps in Coot ^64^, and refined using Phenix ^65^. Structure was visualized using PyMOL ^66^.

## Data Availability

Atomic coordinates and structure factors for the reported TGP-Sb32 structure are deposited in the Protein Data Bank (PDB) under accession codes of 9WPI.

## Acknoledgements

We thank the staff scientists from beamlines BL18U1 at the National Facility for Protein Science in Shanghai (NFPS) at Shanghai Synchrotron Radiation Facility for assistance during the data collection. We thank Prof. Markus Seeger at University of Zurich, Switzerland for providing the sybody libraries.

## Conflict of Interest

The authors declare that they have no conflict of interest.

**Table 1.** Data collection and refinement statistics

| **Sb32-TGP** |  |
| --- | --- |
| **Data collection** |  |
| Space group | *C121* |
| Cell dimensions |  |
| *a*, *b*, *c* (Å) | 162.47, 89.92, 111.24 |
| *α*, *β*, *γ* (°) | 90, 112.12, 90 |
| Wavelength (Å) | 0.97930 |
| Resolution (Å) | 77.19 – 1.82 (1.92 – 1.82)<sup>a</sup> |
| *R*<sub>merge</sub> | 0.086 (0.636) |
| *R*<sub>pim</sub> | 0.043 (0.353) |
| *I*/*σI* | 12.1 (2.6) |
| Completeness (%) | 99.4 (99.3) |
| Multiplicity | 4.9 (4.2) |
| *CC* * <sup>b</sup> | 0.999 (0.960) |
| **Refinement** |  |
| Resolution (Å) | 40.61 – 1.82 |
| No. reflections | 132,462 |
| *R*<sub>work</sub> / *R*<sub>free</sub> | 0.1494/0.1718 |
| No. atoms | 9,234 |
| Protein | 8,007 |
| Chromophore/glycerol | 174 |
| Water | 1,053 |
| No. residues | 997 |
| B-factors (Å<sup>2</sup>) | 36.25 |
| Protein | 35.10 |
| Chromophore/glycerol | 42.80 |
| Water | 43.91 |
| R.m.s deviations |  |
| Bond lengths (Å) | 0.016 |
| Bond angles (°) | 1.37 |
| Ramachandran |  |
| Favoured (%) | 98.67 |
| Allowed (%) | 1.33 |
| Outlier (%) | 0.00 |
| **PDB ID** | **9WPI** |

<sup>a</sup>Highest resolution shell is shown in parenthesis. <sup>b</sup> $CC^* = \sqrt{\frac{2CC_{1/2}}{1+CC_{1/2}}}$

## References

1 Scully, M. et al. Caplacizumab Treatment for Acquired Thrombotic Thrombocytopenic Purpura. N Engl J Med 380, 335–346 (2019).

2 Fan, S., Gai, C., Li, B. & Wang, G. Efficacy and safety of envafolimab in the treatment of advanced dMMR/MSI–H solid tumors: A single–arm meta–analysis. Oncol Lett 26, 351 (2023).

3 Papp, K. A., Weinberg, M. A., Morris, A. & Reich, K. IL17A/F nanobody sonelokimab in patients with plaque psoriasis: a multicentre, randomised, placebo-controlled, phase 2b study. Lancet 397, 1564–1575 (2021).

4 Girard, N., et al. Phase Ib Study of BI 836880 (VEGF/Ang2 nanobody) in Combination With Ezabenlimab (anti–PD-1 antibody) in Patients With Advanced or Metastatic Solid Tumors. JCO Oncology Advances, e2400039 (2025).

5 Fang, Y. et al. Abstract 2857: DR30303, a SMART-VHHBody powered anti-CLDN18.2 VHH-Fc with enhanced ADCC activity for the treatment of gastric and pancreatic cancers. Cancer Res 82, 2857–2857 (2022).

6 Mehra, N. et al. Phase I/II trial of LAVA-1207, a novel bispecific gamma-delta T-cell engager alone, or with low dose IL-2 or pembrolizumab, in metastatic castration resistant prostate cancer (mCRPC). J Clin Oncol 42, TPS2672–TPS2672 (2024).

7 Zhao, Y. et al. KN046, a bispecific antibody against PD-L1 and CTLA-4, plus chemotherapy as first-line treatment for metastatic NSCLC: A multicenter phase 2 trial. Cell Rep Med 5, 101470 (2024).

8 Yi, S. et al. Advancing pancreatic cancer therapy by mesothelin-specific nanobody conjugation. Mol Cancer 24, 124 (2025).

9 Dörner, T. et al. FRI0239 Results of a phase 2b study of vobarilizumab, an anti-interleukin-6 receptor nanobody, as monotherapy in patients with moderate to severe rheumatoid arthritis. Ann Rheum Dis 76, 575 (2017).

10 Cong, Y., Devoogdt, N., Lambin, P., Dubois, L. J. & Yaromina, A. Promising Diagnostic and Therapeutic Approaches Based on VHHs for Cancer Management. Cancers (Basel) 16 (2024).

11 Leeuw, T. et al. Combined TNF-α and OX40L targeting as a new treatment option for hidradenitis suppurativa. J Allergy Clin Immunol Glob 4, 100483 (2025).

12 Cunningham, S. et al. Nebulised ALX-0171 for respiratory syncytial virus lower respiratory tract infection in hospitalised children: a double-blind, randomised, placebo-controlled, phase 2b trial. Lancet Respir Med 9, 21–32 (2021).

13 Rashidian, M. & Ploegh, H. Nanobodies as non-invasive imaging tools. Immunooncol Technol 7, 2–14 (2020).

14 Wang, W. et al. Nanobody-Based PET Imaging of CD47 Expression in Thyroid Carcinoma. Mol Pharm 22, 3414–3422 (2025).

15 Rizk, S. S., Moustafa, D. M., ElBanna, S. A., Nour El-Din, H. T. & Attia, A. S. Nanobodies in the fight against infectious diseases: repurposing nature’s tiny weapons. World J Microbiol Biotechnol 40, 209 (2024).

16 Redrado-Hernández, S. et al. Broad Protection against Invasive Fungal Disease from a Nanobody Targeting the Active Site of Fungal β-1,3-Glucanosyltransferases. Angew Chem Int Ed Engl 63, e202405823 (2024).

17 Li, T. et al. A synthetic nanobody targeting RBD protects hamsters from SARS-CoV-2 infection. Nat Commun 12, 4635 (2021).

18 Yao, H. et al. A high-affinity RBD-targeting nanobody improves fusion partner’s potency against SARS-CoV-2. PLoS Pathog 17, e1009328 (2021).

19 Li, T. et al. Isolation, characterization, and structure-based engineering of a neutralizing nanobody against SARS-CoV-2. Int J Biol Macromol 209, 1379–1388 (2022).

20 Li, T. et al. Structural Characterization of a Neutralizing Nanobody With Broad Activity Against SARS-CoV-2 Variants. Front Microbiol 13, 875840 (2022).

21 Schoof, M., et al. An ultra-potent synthetic nanobody neutralizes SARS-CoV-2 by locking Spike into an inactive conformation. *bioRxiv* (2020).

22 Li, T., Cai, H., Lai, Y., Yao, H. & Li, D. A simple and effective method to remove pigments from heterologous secretory proteins expressed in Pichia pastoris. Adv Biotechnol (Singap) 2, 5 (2024).

23 Wang, Z. et al. ALFA nanobody-guided endogenous labeling. Nat Chem Biol (2025).

24 Martinez-Delgado, G. Inhaled nanobodies against COVID-19. Nat Rev Immunol 20, 593 (2020).

25 Bräuer, P. et al. Structural basis for pH-dependent retrieval of ER proteins from the Golgi by the KDEL receptor. Science 363, 1103–1107 (2019).

26 Guo, X. et al. Structure and mechanism of human cystine exporter cystinosin. Cell 185, 3739–3752.e3718 (2022).

27 Löbel, M. et al. Structural basis for proton coupled cystine transport by cystinosin. Nat Commun 13, 4845 (2022).

28 Kumar, H. et al. Crystal Structure of a ligand-bound LacY-Nanobody Complex. Proc Natl Acad Sci U S A 115, 8769–8774 (2018).

29 Yang, Z. et al. Structural insights into auxin recognition and efflux by Arabidopsis PIN1. Nature 609, 611–615 (2022).

30 Brunner, J. D. & Schenck, S. Production and Application of Nanobodies for Membrane Protein Structural Biology. Methods Mol Biol 2127, 167–184 (2020).

31 Uchański, T. et al. Megabodies expand the nanobody toolkit for protein structure determination by single-particle cryo-EM. Nat Methods 18, 60–68 (2021).

32 Coupland, C. E. et al. Structure, mechanism, and inhibition of Hedgehog acyltransferase. Mol Cell 81, 5025–5038.e5010 (2021).

33 Bloch, J. S. et al. Development of a universal nanobody-binding Fab module for fiducial-assisted cryo-EM studies of membrane proteins. Proc Natl Acad Sci U S A 118, e2115435118 (2021).

34 Wu, X. & Rapoport, T. A. Cryo-EM structure determination of small proteins by nanobody-binding scaffolds (Legobodies). Proc Natl Acad Sci U S A 118, e2115001118 (2021).

35 Kang, Y. & Chen, L. Structural basis for the binding of DNP and purine nucleotides onto UCP1. Nature 620, 226–231 (2023).

36 Lin, H. et al. Structure and mechanism of the plastid/parasite ATP/ADP translocator. Nature 641, 797–804 (2025).

37 Yi, G. et al. Covalently constrained ‘Di-Gembodies’ enable parallel structure solutions by cryo-EM. Nat Chem Biol (2025).

38 Kruse, A. C. et al. Activation and allosteric modulation of a muscarinic acetylcholine receptor. Nature 504, 101–106 (2013).

39 Hillier, J. et al. Structural insights into Frizzled3 through nanobody modulators. Nat Commun 15, 7228 (2024).

40 Rasmussen, S. G. et al. Structure of a nanobody-stabilized active state of the β(2) adrenoceptor. Nature 469, 175–180 (2011).

41 Rasmussen, S. G. et al. Crystal structure of the β2 adrenergic receptor-Gs protein complex. Nature 477, 549–555 (2011).

42 Song, A. & Wu, X. Mechanistic insights of substrate transport and inhibitor binding revealed by high-resolution structures of human norepinephrine transporter. Cell Res 34, 810–813 (2024).

43 Irobalieva, R. N. et al. Structural Basis of the Allosteric Inhibition of Human ABCG2 by Nanobodies. J Mol Biol 435, 168234 (2023).

44 Close, D. W. et al. Thermal green protein, an extremely stable, nonaggregating fluorescent protein created by structure-guided surface engineering. *Proteins: Structure*, Function, and Bioinformatics 83, 1225–1237 (2015).

45 Xu, Y. et al. Structures of liganded glycosylphosphatidylinositol transamidase illuminate GPI-AP biogenesis. Nat Commun 14, 5520 (2023).

46 Xu, Y. et al. Molecular insights into biogenesis of glycosylphosphatidylinositol anchor proteins. Nat Commun 13, 2617 (2022).

47 Hong, J. et al. Molecular basis of the inositol deacylase PGAP1 involved in quality control of GPI-AP biogenesis. Nat Commun 15, 8 (2024).

48 Xiong, Q. et al. Molecular architecture of human LYCHOS involved in lysosomal cholesterol signaling. Nat Struct Mol Biol 32, 905–913 (2025).

49 Cai, H. et al. An improved fluorescent tag and its nanobodies for membrane protein expression, stability assay, and purification. Commun Biol 3, 753 (2020).

50 Yao, H., Cai, H. & Li, D. Fluorescence-Detection Size-Exclusion Chromatography-Based Thermostability Assay for Membrane Proteins. Methods Mol Biol 2564, 299–315 (2023).

51 Hattori, M., Hibbs, R. E. & Gouaux, E. A fluorescence-detection size-exclusion chromatography-based thermostability assay for membrane protein precrystallization screening. Structure 20, 1293–1299 (2012).

52 Yue, Z. et al. Structure-based design of covalent nanobody binders for a thermostable green fluorescence protein. Acta Biochim Biophys Sin (Shanghai) 57, 1363–1370 (2024).

53 Zimmermann, I. et al. Synthetic single domain antibodies for the conformational trapping of membrane proteins. eLife 7, e34317 (2018).

54 Myung, Y., Pires, D. E. V. & Ascher, D. B. Understanding the complementarity and plasticity of antibody-antigen interfaces. Bioinformatics 39 (2023).

55 Kourelis, J., Marchal, C., Posbeyikian, A., Harant, A. & Kamoun, S. NLR immune receptor–nanobody fusions confer plant disease resistance. Science 379, 934–939 (2023).

56 Kirchhofer, A. et al. Modulation of protein properties in living cells using nanobodies. Nat Struct Mol Biol 17, 133–138 (2010).

57 Zhang, Z., Wang, Y., Ding, Y. & Hattori, M. Structure-based engineering of anti-GFP nanobody tandems as ultra-high-affinity reagents for purification. Sci Rep 10, 6239 (2020).

58 Li, T. et al. A synthetic nanobody targeting RBD protects hamsters from SARS-CoV-2 infection. Nat Commun 12, 4635 (2021).

59 Cai, H., Yao, H., Li, T., Tang, Y. & Li, D. High-level heterologous expression of the human transmembrane sterol Δ8,Δ7-isomerase in Pichia pastoris. Protein Expr Purif 164, 105463 (2019).

60 Cai, H. et al. An improved fluorescent tag and its nanobodies for membrane protein expression, stability assay, and purification. Commun Biol 3, 753 (2020).

61 Kabsch, W. XDS. Acta Crystallogr. D Biol. Crystallogr. 66, 125–132 (2010).

62 Evans, P. R. & Murshudov, G. N. How good are my data and what is the resolution? Acta Crystallogr. D Biol. Crystallogr. 69, 1204–1214 (2013).

63 McCoy, A. J. et al. Phaser crystallographic software. J. Appl. Crystallogr. 40, 658–674 (2007).

64 Emsley, P., Lohkamp, B., Scott, W. G. & Cowtan, K. Features and development of Coot. Acta Crystallogr. D Biol. Crystallogr. 66, 486–501 (2010).

65 Adams, P. D. et al. PHENIX: a comprehensive Python-based system for macromolecular structure solution. Acta Crystallogr. D Biol. Crystallogr. 66, 213–221 (2010).

66 Schrödinger, L. L. C. *The PyMOL Molecular Graphics System, Version 1.8*, 2015).

